# Whole Genome Sequencing and Rare Variant Analysis in Essential Tremor Families

**DOI:** 10.1101/248443

**Authors:** Zagaa Odgerel, Nora Hernandez, Jemin Park, Ruth Ottman, Elan D. Louis, Lorraine N. Clark

## Abstract

Essential tremor (ET) is one of the most common movement disorders. The etiology of ET remains largely unexplained. Whole genome sequencing (WGS) is likely to be of value in understanding a large proportion of ET with Mendelian and complex disease inheritance patterns. In ET families with Mendelian inheritance patterns, WGS may lead to gene identification where WES analysis failed to identify the causative variant due to incomplete coverage of the entire coding region of the genome. Alternatively, in ET families with complex disease inheritance patterns with gene x gene and gene x environment interactions enrichment of functional rare coding and non-coding variants may explain the heritability of ET. We performed WGS in eight ET families (n=40 individuals) enrolled in the Family Study of Essential Tremor. The analysis included filtering WGS data based on allele frequency in population databases, rare variant classification and association testing using the Mixed-Model Kernel Based Adaptive Cluster (MM-KBAC) test and prioritization of candidate genes identified within families using phenolyzer. WGS analysis identified candidate genes for ET in 5/8 (62.5%) of the families analyzed. WES analysis in a subset of these families in our previously published study failed to identify candidate genes. In one family, we identified a deleterious and damaging variant (c.1367G>A, p.(Arg456Gln)) in the candidate gene, *CACNA1G*, which encodes the pore forming subunit of T-type Ca(2+) channels, Ca_V_3.1, and is expressed in various motor pathways and has been previously implicated in neuronal autorhythmicity and ET. Other candidate genes identified include *SLIT3* (family D), which encodes an axon guidance molecule and in three families, phenolyzer prioritized genes that are associated with hereditary neuropathies (family A, *KARS*, family B, *KIF5A* and family F, *NTRK1*). This work has identified candidate genes and pathways for ET that can now be prioritized for functional studies.

## INTRODUCTION

Essential tremor (ET) is one of the most common neurological disorders. In most studies the prevalence of ET is markedly higher than that of Parkinson’s disease (PD). The prevalence of ET is estimated to be 2.2% and as much as 4.6% in cases aged ≥65 years [1]. The defining clinical feature of ET is a kinetic tremor at 4-12 Hz. This tremor occurs in the arms and hands; it may also eventually spread to involve several cranial regions (e.g., the neck, voice, and jaw). Both genetic and environmental (i.e., toxic) factors are likely to contribute to the etiology of ET. The high heritability and aggregation of ET in families suggests a Mendelian pattern of inheritance [2-5]. Family studies indicate that on the order of 30 - 70% of ET patients have a family history with the vast majority (>80%) of young-onset (<40 years old) cases reporting >1 affected first-degree relative [6].

Four published genome wide linkage scans have been performed all in North American or Icelandic ET families [7-9]. These studies led to the identification of genetic loci harboring ET genes on chromosomes 3q13 (ETM1 OMIM:190300) [7], 2p22-p25 (ETM2 OMIM:602134) [8], 6p23 (ETM3 OMIM: 611456) [9], and 5q35 [10]. Recently, several studies have used a whole exome sequencing (WES) approach to identify candidate genes in ET families [11-16]. Collectively, these studies suggest that ET is genetically heterogeneous.

With the limited nature of this progress, the genetic etiology of ET still remains largely unexplained. Whole genome sequencing (WGS) is likely to be of value in furthering our understanding of a large proportion of ET where WES analysis has failed to identify the causative variant [17]. WGS which forgoes capturing is less sensitive to GC content and is more likely than WES to provide complete coverage of the entire coding region of the genome [18].

Here we report analysis of eight early-onset ET families (n=40 individuals) enrolled in the family study of Essential Tremor (FASET) at Columbia University. The analysis included filtering on WGS data based on allele frequency in population databases, rare variant classification and association using the Mixed-Model Kernel Based Adaptive Cluster (MM-KBAC) test [19, 20], and prioritization of candidate genes identified within families using phenolyzer.

## MATERIALS AND METHODS

### Study participants and clinical diagnosis

Study subjects and relatives were enrolled in a family study of ET at Columbia University NY, USA. The study was approved by the Institutional Review Board at Columbia University and written informed consent was obtained from all participants. Details of the study, criteria for enrollment, and diagnosis of ET has been described previously [15].We selected a total of 8 families for WGS (n=40 individuals), which included affected and unaffected first-degree relatives. The eight families have been previously described in a WES study [15]. All affected individuals included in the study received a diagnosis of definite, probable or possible ET. Possible and probable ET family members were considered affected. The criteria we used, namely, the Washington Heights Inwood Genetic Study of ET (WHIGET) criteria are very strict [21]. All ET diagnoses (possible, probable and definite) required, at a minimum, moderate or greater amplitude kinetic tremor on at least three tasks, and an absence of other etiologies. As such, these criteria for all three categories of ET (i.e., possible, probable and definite) are even more stringent than those for definite ET that were outlined in the original Consensus Statement on Tremor of the Movement Disorders Society (published in 1998) [22] and the revised Consensus Criteria (published in 2017) [23]. The clinical characteristics of study participants are summarized in Table 1 and pedigrees of the families are shown in Fig 1.

**Table 1.** Clinical characteristics of affected ET individuals and unaffected family members that were whole genome sequenced in eight families

| Clinical characteristic | ET patients<br>n=31 | Unaffected<br>n=9 | Total<br>n=40 |
| --- | --- | --- | --- |
| Male n (%) | 12 (39) | 3 (33) | 15 (38) |
| Age at tremor onset mean years (SD) | 27.83 (19.30) | NA | NA |
| Age at interview mean years, SD | 58 (18.08) | 56.63 (13.65) | 57.72 (17.11) |
| Duration of tremor mean years, SD | 30.47 (18.98) | NA | NA |
| Total tremor score mean SD | 17.76±7.80 (39) | NA | NA |
| Head tremor on examination n (%) | 12 (39) | NA | NA |
| Chin tremor on examination n (%) | 6 (19) | NA | NA |
| Head tremor presence in head | 4 (13) | NA | NA |
| and chin n (%) |  |  |  |

**Figure 1.**
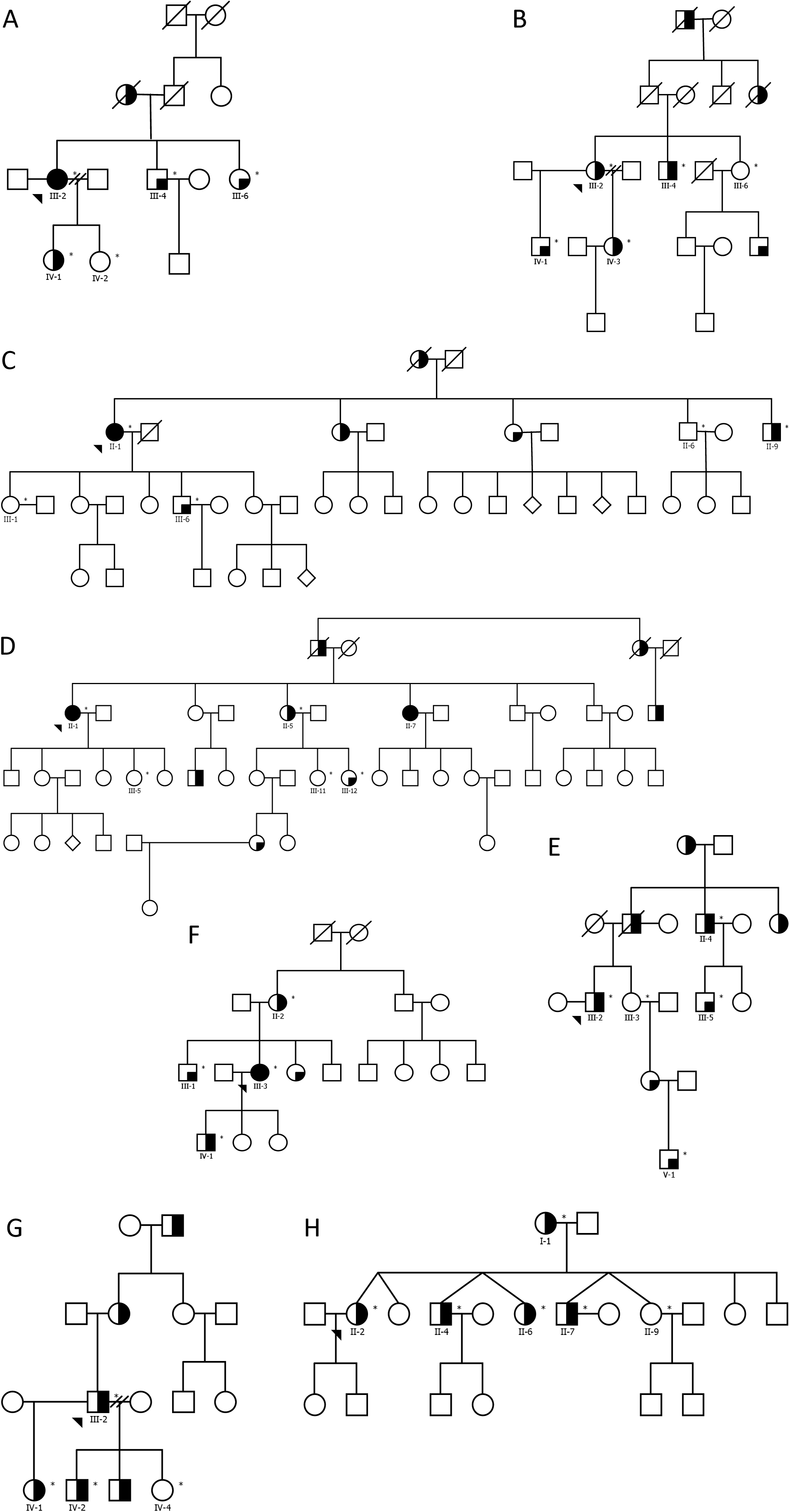
Pedigrees of eight ET families that were whole genome sequenced. Pedigrees for families (A-H) that were whole genome sequenced are shown. The generation in each pedigree is shown by roman numerals. The proband is indicated by an arrowhead. A^‘*’^ symbol indicates subjects that were whole genome sequenced. Below each subject with DNA avaliable for genetic analysis the subject ID is indicated. Symbol shading is as follows: definite ET, symbols completely black; probable ET, symbols half vertical black fill; possible ET, symbols with a quadrant in black; and unaffected clear symbol. To protect the identity of participants in families the gender and birth order were changed in order to disguise their identities.

### Whole Genome Sequencing and quality control

Genomic DNA was isolated from peripheral blood cells using standard methods. Whole genome sequencing was performed on the genomic DNA of 4-5 individuals including affected and unaffected (definite, probable or possible ET diagnosis) individuals from each of eight families. The pedigrees of eight families are shown in Fig.1. Libraries were prepared using the TruSeq DNA PCR-free kit (Illumina San Diego CA USA). Paired-end sequencing (2x150 bp) was performed at >30x coverage per sample. Resulting libraries were sequenced on Illumina HiSeq TENx (Illuminia San Diego CA). Sequence alignment to the UCSC hg19 reference genome was performed using the Burrows-Wheeler Aligner algorithm [24] and variant calling was performed using the Genome Analysis Toolkit (GATK; Broad Institute Cambridge MA USA) [25]. Duplicate reads were removed using Picard (http://broadinstitute.github.io/picard/). Local realignment and quality recalibration was performed via GATK. Quality control checks for samples were performed according to GATK best practices.

### Variant filtering based on allele frequency in population databases

We filtered and removed variants with MAF≥0.001 in all individuals in 1000 Genomes Phase 3 or the NCBI dbSNP common 147 database, resulting in a total of 3,777,271 rare variants across all samples.

#### Classification of rare variants based on variant type

Assessment of variants was performed based on reference sequence GRCh37 and RefSeq Gene transcripts of NBCI Homo sapiens Annotation Release 105 that was implemented in the Golden Helix SNP & Variation Suite (SVS) ver.8.2 (Golden Helix MT). Rare variants were classified into five groups, based on localization to a gene region and predicted effect on transcript and protein: 1) 5’-UTR and 3’-UTR (n=26,872 variants in 8,299 genes), 2) nonsynonymous (n=11,272 variants in 4,877 genes), 3) loss-of-function (LoF) (n=1,365 variants in 711 genes), 4) synonymous (n=5,854 variants in 3,164 genes), and 5) intronic (n=1,174,082 variants in 16,486 genes). LoF variants were defined as follows: nonsense variants that introduce stop gain/loss of codons, variants that disrupt splice sites including canonical splice donor and acceptor sites and frameshift variants that disrupt a transcript’s open reading frame.

### Annotation of Variants

Rare variants were assessed using several *in silico* tools including the Combined Annotation Dependent Depletion (CADD) tool [26] implemented in the Golden Helix SNP & Variation Suite (SVS) ver.8.6.0 (Golden Helix MT). CADD measures deleteriousness of variants (coding and non-coding intronic) that is a property strongly correlated with molecular functionality and pathogenicity [27]. Variants were filtered based on a phred-scaled CADD score and variants with a phred-scaled CADD score>10, corresponding to the top 10% of deleterious substitutions relative to all possible variants in the human reference genome [26] were retained for further analyses. We also assessed deleteriousness of variants using several *in silico* tools including SIFT [28], PolyPhen2 [29], LRT [30], Mutation Taster [31], FATHMM [32], PROVEAN [33], MetaSVM and MetaLR [34] as implemented in the Golden Helix SNP & Variation Suite (SVS) ver.8.6.0 (Golden Helix MT). Only variants with a phred-scaled CADD score>10 and/or predicted to be deleterious or damaging by ≥1 *in silico* prediction tool were retained for further analysis.

#### Synonymos variants in splicing regulatory regions

To determine whether synonymous variants identified in our analyses are enriched in splicing enhancer regions and splicing silencer regions we used http://genes.mit.edu/burgelab/rescue-ese/ and http://genes.mit.edu/fas-ess/ online tools, respectively [35,36].

#### Non-coding intronic variants in DNase I hypersensitivity and transcription factor binding sites

We performed further evaluation of non-coding intronic variants by assessing whether these variants are enriched in *DNase* I hypersensitive sites that represent open chromatin regions accessible to transcription factors. We downloaded the wgEncodeRegDnaseClusteredV3 table from the DNAse Clusters track which contains DNaseI Hypersensitive Sites in 125 cell types in ENCODE (http://genome.ucsc.edu/cgi-bin/hgTables) [37].

#### Residual Variation Intolerance Score (RVIS)

We assessed the candidate genes identified in this study to determine whether they are intolerant to variants by applying the residual variation intolerance score (RVIS) [38].

### MM-KBAC Analysis

We performed a rare variant classification and association analysis using the regression and permutation based Mixed-Model Kernel-Based Adaptive Cluster method (MM-KBAC) [19], and the within gene interaction model to analyze rare functional variants, as implemented in SVS ver.8.6.0 (Golden Helix MT). KBAC catalogs rare variant data within a gene region/transcript (genome-wide) into multi-marker genotypes and determines their association with the phenotype, weighing each multi-marker genotype by how often that genotype was expected to occur according to control and case data and the null hypothesis that there is no association between the genotype and the case/control status. Thus, genotypes with high sample risks are given higher weights that potentially separate causal from non-causal genotypes. The logistic mixed model approach for KBAC to adjust for family structure and relatedness was used and has been described previously [20]. Possible and probable ET family members were considered affected. The control population used included unaffected family members. A *p* value was assessed by an adaptive permutation procedure in association tests [19]. The test applied 10 000 permutations and an adaptive permutation threshold of alpha 0.01 and used the earliest start position and the last stop position of all transcripts to define a gene based on the RefSeq Gene transcripts 105v2 NCBI. By default, variants flanking (proximal and distal) the gene region up to a distance of 1000 bp were included in the analysis. We selected genes with a *p* value<0.05 for further analysis. The analysis was performed separately for variants classified by variant type in the dataset. When MM-KBAC analysis was performed separately for variants based on variant type (nonsynonymous, LoF, 5’UTR and 3’UTR, synonymous and intronic) the total number of genes with *p* value <0.05 was 163.

### Co-segregation of variants with ET within families

Variants identified from the MM-KBAC analysis, that were annotated with a phred scaled score>10 by CADD (coding and non-coding intronic variants) and/or predicted by *in silico* prediction tools to be deleterious or damaging (coding variants) were assessed for co-segregation with ET within families. The criteria that we used to define co-segregation is as follows: 1) the annotated variant was present in all affected ET individuals and 2) absent from unaffected individuals within a family. Sanger sequencing was used to validate and confirm variants within a family and to genotype family members with available DNA that did not have WGS data. Genes harboring variants that were annotated with a phred scaled score>10 by CADD (coding and non-coding intronic variants) and/or predicted by *in silico* prediction tools to be deleterious or damaging (coding variants) and that co-segregated with ET within single family were prioritized for phenolyzer.

### Prioritization of Candidate Genes using Phenolyzer

Phenolyzer is a computational tool that uses prior information to implicate genes involved in diseases [39]. Phenolyzer exhibits superior performance over competing methods for prioritizing Mendelian and complex disease genes based on disease or phenotype terms entered as free text. The most disease relevant genes, considering all reported gene-disease relationships, are shown as seed genes. Predicted genes are input (seed) genes that are expanded to include related genes on the basis of several gene-gene relationships (e.g. protein-protein interactions, biological pathway, gene family or transcriptional regulation). The following disease/phenotype terms were used: Tremor, Essential Tremor, Parkinson’s disease, Channelopathy, Epilepsy, neurological, neurodegenerative, Spinocerebellar ataxia, Fragile X Associated Tremor Ataxia Syndrome, brain, cerebellar diseases. For each family, candidate genes with prioritized variants were uploaded as input for phenolyzer analysis. The gene disease score and gene prediction score system is described in Yang et al., 2015 [39]. Phenolyzer generates raw and normalized scores for seed and predicted genes [39].

### Availability of Data

All phenotype and whole genome sequence data generated from this study will be released and deposited in the database of Genotypes and Phenotypes (dbGaP; http://www.nlm.nih.gov/gap) of the National Center for Biotechnology Information. The study titled ‘Identification of Susceptibility Genes for Essential Tremor’ received the dbGaP Study Accession: phs000966.v1.p1. Additionally, all deidentified WGS data and related meta data underlying the findings reported in this manuscript will be made available at the public repository Dryad (datadryad.org).

## RESULTS

To identify candidate genes in ET we conducted WGS in 40 individuals from 8 families with multiple affected ET members (Table 1 and Fig.1). Datasets were generated based on filtering of variants on allele frequency in population databases (Fig. 2). To identify and prioritize genes in the ET families we performed rare variant classification and association analysis using the Mixed-Model Kernel Based Adaptive Cluster (MM-KBAC) test [19] followed by phenolyzer [39].

**Figure 2.**
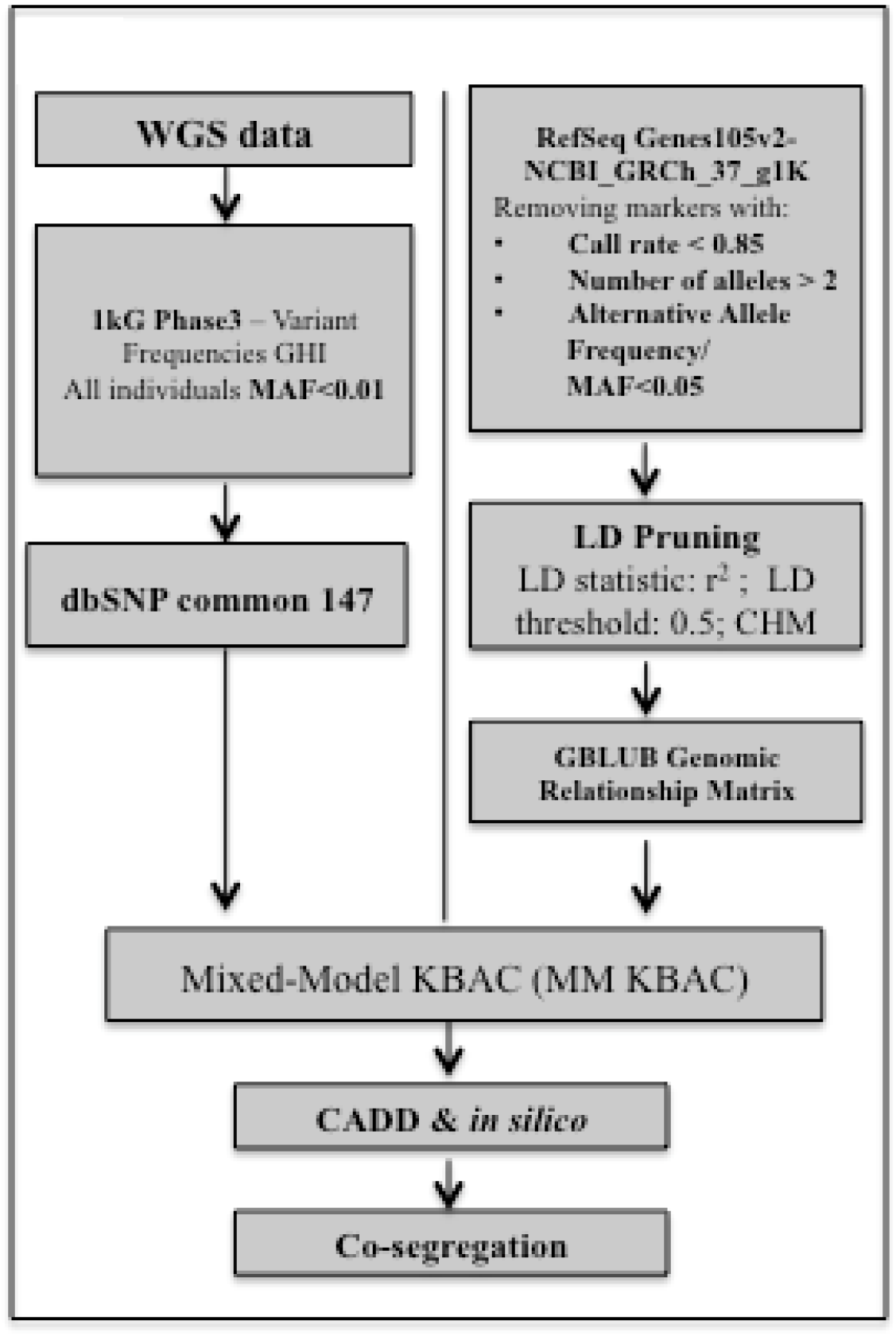
Analysis workflow for analysis using MM-KBAC. The analysis workflow for WGS data is shown with population database filtering, analysis methods and annotation.

### Rare Variant Classification and Association Analysis of rare variants with MAF≤0.01

After QC and variant filtering, a total of 3,777,271 variants were selected for the subsequent analyses (Fig. 2). By MM-KBAC analysis, we obtained 1,325 genes with *p* value<0.05 (with-in gene association) and 3,779 variants located within these genes. Of those, 783 variants were annotated with a phred scaled score>10 by CADD and 95 variants were predicted by *in silico* prediction tools to be deleterious or damaging. We assessed the following variant types: 1) nonsynonymous, 2) LoF, 3) 5’UTR and 3’UTR, 4) synonymous and 5) intronic variants. Variants identified from the MM-KBAC analysis, that were annotated with a phred scaled score>10 by CADD and/or predicted by *in silico* prediction tools to be deleterious or damaging and that co-segregated within the ET families are shown in Table 2. A total of 168 variants located in 163 genes co-segregated with ET within families.

**Table 2.**
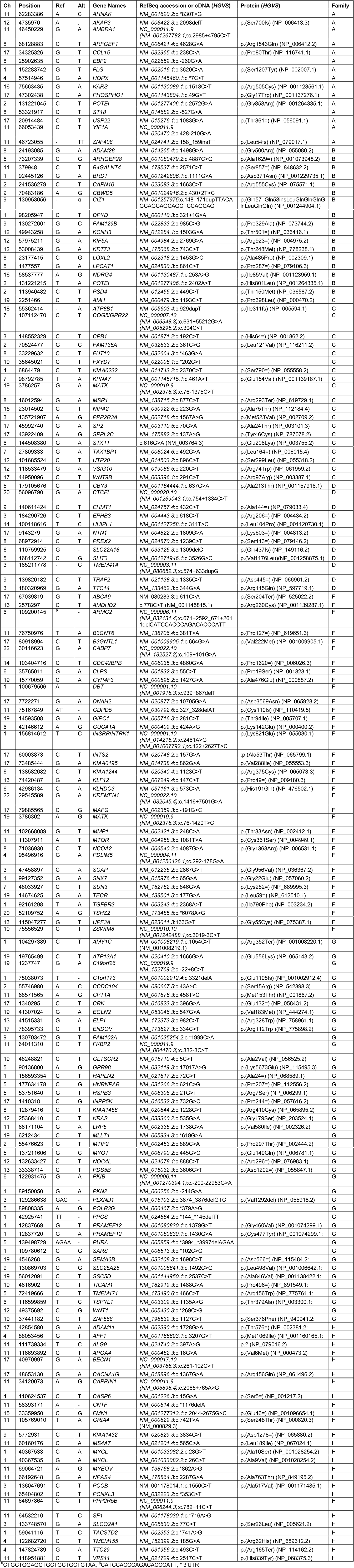
Variants identified in families co-segregating wit ET based on MM-KBAC analysis of rare variants by variant type

#### Nonsynonymous variants

We conducted the MM-KBAC analysis on 11,272 rare nonsynonymous variants in 4,877 genes and obtained a total of 316 genes with *p*<0.05. After annotation of variants, we obtained 87 variants that co-segregated within families. One variant in *COPZ2* was removed from analysis based on the MAF>0.01 reported in ExAC although it was not reported in the 1000 Genomes data.

#### LoF variants

The analysis was performed on 1,364 rare LoF variants located in 711 genes and a total of 60 genes were obtained with a *p*<0.05. Following annotation, 13 deleterious variants co-segregated within families (Table 2). For Indel variants, BAM files were manually examined using the Genome browser in SVS v8.6 (Golden Helix) to verify the variant.

#### Variants in 5’-UTR and 3-UTR regions

The MM-KBAC analysis was conducted on 26,872 rare variants in 8,299 genes and 409 genes were obtained with a *p*<0.05. Following annotation of variants and analysis of co-segregation, 25 variants co-segregated within families (Table 2).

#### Synonymous variants

The analysis was performed on 5,854 synonymous rare variants located in 3,164 genes and a total of 216 genes with a *p* value<0.05 were obtained. Following annotation, a total of 35 variants co-segregated within families (Table 2). A variant in *ASB16* was excluded from the analysis based on the allele frequency reported in ExAC (MAF=0.0278). We also investigated whether synonymous variants were located in splicing enhancer and silencer regions within genes. The variants c.429G>A (NM_006024.6), c.3606C>T, c.1809G>A and c.177G>A were identified in enhancer regions in the *TAX1BP1, PDS5B*, *NTN1* and *TECR* genes respectively and c.72C>T (NM_02817.2), c.846G>A (NM_152782.3), and c.861C>T (NM_024830.3) were located in splicing silencer regions in the *HAPLN2, SUN3*, and *LPCAT1* genes respectively (Table 3).

**Table 3.**
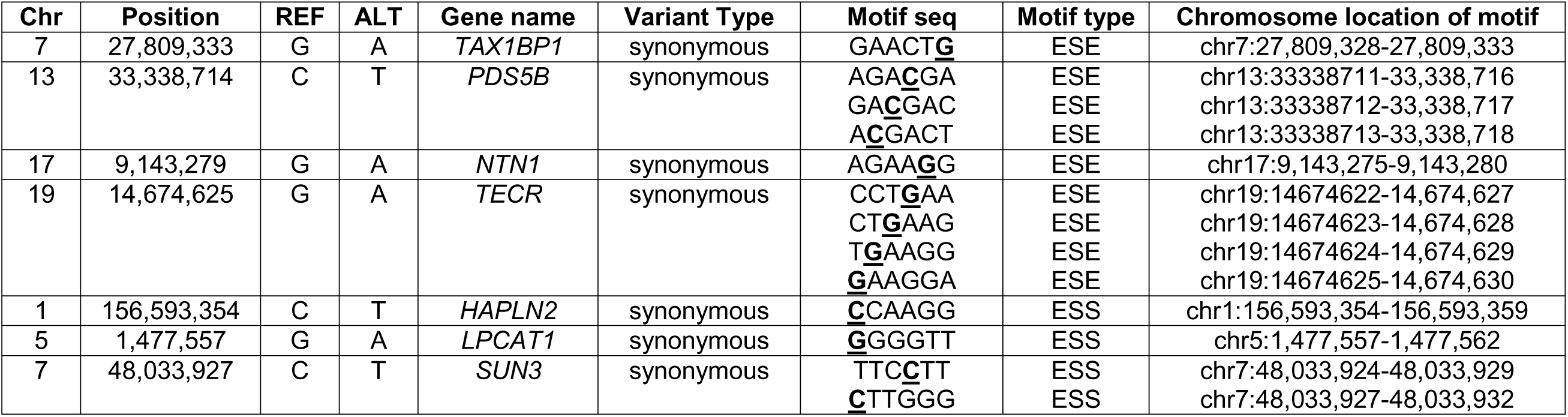
Synonymous variants in enhancer and splicing regions identified in families co-segregating with ET based on MM-KBAC analysis of rare variants

#### Intronic variants

The MM-KBAC analysis was conducted on 1,174,082 intronic rare variants located in 16,486 genes and 324 genes with a *p* value<0.05 were obtained. Following annotation and co-segregation analysis, we obtained a total of 14 deleterious variants that co-segregated within families (Table 2).

#### DNAse I Hypersensitivity Sites and Transcription Factor Binding Sites

Genetic variants can affect transcription factor binding sites (TFBS), particularly via their enrichment in *DNase I* hypersensitive sites (DHS) that provide open chromatin access to transcription factors. Thus we sought variants that could be enriched at these sites using TFBS conserved data in ENCODE [40]. We asked whether the 169 variants (MM-KBAC analysis by variant type, and that includes annotated variants that co-segregated within ET families) identified from our analyses were found in DHS. 67 variants in 65 genes were in DHS. These 67 variants comprised 6 of 67 (9%) 5’-UTR variants; 6 of 67 (9%) 3’-UTR variants; 3 of 67 (4%) were LoF variants; 36 of 67 (54%) were non-synonymous variants; 12 of 67 (18%) were synonymous; and 4 of 67 (6%) intronic variants. DHSs are enriched with transcription factor binding sites (TFBSs), crucial sequences for the regulation of gene expression. Cross species conservation of genomic sequence has been successfully used for identifying biologically functional TFBS [41]. We identified 40 variants within TFBS (Table 4).

**Table 4.** Variants located within TFBS identified in families co-segregating with ET

| Chromosome | Position | Reference | Alternates | Transfac binding matrix id | Strand | Family |
| --- | --- | --- | --- | --- | --- | --- |
| 1 | 11307911 | A | T | *TCF11MAFG_01* | + | F |
| 1 | 40367533 | C | A | *ELK1_01* | + | H |
| 1 | 40367535 | G | A | *ELK1_01* | + | H |
| 1 | 92161298 | T | A | *CART1_01* | - | F |
| 2 | 70524477 | G | C | CREB_02 | + | C |
| 3 | 47458897 | C | A | MAZR_01 | + | F |
| 3 | 135721907 | A | G | YY1_01 | - | C |
| 3 | 136047691 | C | T | LUN1_01 | + | H |
| 4 | 88053456 | G | T | YY1_01 | - | H |
| 4 | 95496916 | G | A | PAX4_04 | + | F |
| 5 | 53751640 | G | T | HTF_01 | + | G |
| 5 | 72419666 | C | T | SEF1_C | - | G |
| 5 | 90136800 | A | G | MEF2_04 | + | G |
| 6 | 42146612 | A | G | COMP1_01 | + | F |
| 6 | 42986134 | C | A | HOX13_01 | + | F |
| 6 | 122931475 | G | A | SP1_Q6 | + | G |
| 8 | 23177415 | C | G | AHRARNT_02 | + | B |
| 8 | 53321917 | C | T | AREB6_01 | - | A |
| 8 | 71036930 | C | T | AREB6_04 | + | F |
| 11 | 60160176 | C | A | NRSF_01 | - | H |
| 11 | 62283386 | A | C | HNF1_01 | + | A |
| 11 | 66192648 | G | A | AREB6_04 | - | H |
| 11 | 68171104 | G | A | TCF11_01 | + | G |
| 11 | 69064721 | A | G | TBP_01 | + | H |
| 11 | 75167849 | AT | - | PPARA_01 | - | F |
| 11 | 102668089 | G | T | AREB6_04 | + | F |
| 13 | 74420487 | G | A | SRF_01 | - | F |
| 14 | 103404716 | C | T | P53_01 | + | F |
| 17 | 1410318 | C | G | PAX3_01 | - | G |
| 17 | 7722271 | G | A | CREB_02 | + | F |
| 17 | 42854580 | G | A | PAX5_01 | + | H |
| 17 | 43922409 | A | G | TAXCREB_01 | - | C |
| 17 | 73485444 | G | A | NRSF_01 | - | F |
| 17 | 79885565 | C | G | AP4_01 | - | F |
| 17 | 80918994 | C | T | PAX4_01 | - | F |
| 18 | 55362414 | - | A | TCF11_01 | - | C |
| 19 | 1237747 | G | A | PAX5_01 | - | G |
| 19 | 4816902 | C | T | HEN1_01 | + | G |
| 19 | 19765499 | C | T | PPARA_01 | - | G |
| 19 | 48248821 | C | T | YY1_02 | - | G |

### Phenolyzer Analysis

We used phenolyzer to prioritize candidate genes within ET families. The results of the phenolyzer network analysis for 5 families (A, B, D, F, H) are shown in S1 Fig.

### Family A

*KARS* is predicted to be the most disease relevant seed gene (raw score 0.03532; normalized score 0.004) because it maps to Charcot Marie Tooth disease recessive intermediate b in OMIM (OMIM 613641) and DISGENET (C3150897)(S1 Fig.). The nonsynonymous variant identified in *KARS* (c.1513C>T (NM_001130089.1), p.(Arg505Cys)) has a Phred scaled CADD score of 28.6 and is predicted to be deleterious or damaging by several *In Silico* tools (SIFT, POLYPHEN2, Mutation Taster, FATHMM, Provean, MetaSVM and Meta LR). The top three predicted genes are *ARGEF1* (normalized score 0.011), *PHOSPHO1* (normalized score 0.008) and *AMBRA1* (normalized score 0.004).

### Family B

*KIF5A* is predicted to be the most disease relevant seed gene (raw score 0.2954; normalized score 0.033) because it maps to spastic paraplegia 10 in OMIM (OMIM 604187) and DISGENET (C1858712). The variant identified in *KIF5A* is a synonymous variant (c.2769G>A (NM_004984.2), p.(Arg923=)) with a phred-scaled CADD score of 10.95. The nucleotide c.2769 (NM_004984.2) (Chr12:57,975,211) is highly evolutionarily conserved and the FAS-ESS web tool identifies the exon splicing motif ‘CCACTA’ in close proximity (Chr12:57,975,217-57,975,222). The top four predicted genes include *ARHGEF28* (raw score 0.1506; normalized score 0.016), *PSD4* (raw score 0.1208; normalized score 0.013), *LPCAT1* (raw score 0.09227; normalized score 0.01) and *KCNH3* (raw score 0.08023; normalized score 0.008) based on their protein interactions, the same biosystem (e.g. *ARHGEF28*, biosystem Axon guidance, EH-Ephrin signaling and developmental biology), the same gene family (e.g. *PSD4;* gene family, Pleckstrin homology (PH) domain containing or *KCNH3;* gene family, potassium channels Voltage-gated ion channels) or transcription interactions (e.g. *LPCAT1* regulated by ETS1 transcription factor).

### Family C

The top ranked gene is a predicted gene, *MATK* (raw score 0.4266; normalized score 0.046) based on protein interactions (e.g. yeast 2-hybrid with *EWSR1*), the same biosystem (e.g. signal transduction, neurotrophic factor-mediated Trk receptor signalling), the same gene family (e.g. SH2 domain containing) or transcription interactions (e.g. regulated by *GATA2).* The next top three genes (predicted) are *WNT9B* (normalized score 0.025), *TAX1BP1* (normalized score 0.015) and *PPP2R3A* (normalized score 0.015).

### Family D

*SLIT3* is predicted to be one of the most disease relevant seed gene, with a raw score of 0.1637 and normalized score of 0.017, respectively (S1 Fig.). *SLIT3* maps to temporal lobe epilepsy in DISGENET (C0014556) but a disease association with *SLIT3* has not been described in OMIM. The non-synonymous variant identified in *SLIT3* (c.3505G>C (NM_001271946.1), p.(Val1169Leu)); rs144799628) has a Phred scaled CADD score of 22.5 and is predicted to be deleterious or damaging by several *in silico* tools (LRT Pred, Mutation Taster, and FATHMM). The top three predicted genes are *TRAF2* (normalized score 0.035), *EPHB3* (normalized score 0.016) and *SLC22A16* (normalized score 0.01). The variants identified in *TRAF2* (c.1335C>T (NM_021138.3), p.(Asp445=)); phred scaled CADD score of 10.96) and *EPHB3* (c.618C>T (NM_004443.3), p.(Arg206=)); phred scaled score 13.71) are synonymous variants with weak evidence for pathogenicity. The *SLC22A16* (also known as *OCT6*) variant (c.1309delC (NM_033125.3), p.(Gln437fs)) is a LoF frameshift variant, with a phred-scaled CADD score of 35, that is predicted to result in premature termination of the protein.

### Family E

No annotated (phred-scaled CADD score >10 or predicted deleterious or damaging by *in silico* tools) segregating rare deleterious variants were identified in Family E

### Family F

The top predicted disease relevant seed gene is *NTRK1* (raw score 5.152; normalized score 0.538) based on disease mapping to congenital sensory neuropathy with anhidrosis, hereditary sensory and autonomic neuropathy IV (HSAN4) and familial dysautonomia type II in OMIM (OMIM 256800), DISGENET (C0020074), and ORPHANET (642). The variant identified in *NTRK1* is an intronic variant (intron 2;NM_001007792.1:c.122+2627T>C) located in an ENCODE annotated open chromatin/TFBS region (openChrom_2127) of the *NTRK1* gene. The top three predicted genes are *GIPC1* (normalized score 0.06), *MATK* (0.045) and *NCOA2* (normalized score 0.04).

### Family G

The top ranked and predicted seed gene is *CRK* (raw score 0.6991; normalized score 0.073) based on disease mapping in DISGENET, protein interactions (PUBMED 16713569; yeast 2-hybrid with *ATXN1*, score 0.004856), the same biosystem (e.g. signal transduction; *NGF* signaling via *TRKA* form the plasma membrane; signal transduction; signalling to ERKs; signalling by *NGF;* neurotrophic factor-mediated Trk receptor signaling), the same gene family (e.g. SH2 domain containing) or transcription interactions.

### Family H

*CACNA1G* is predicted to be the most disease relevant seed gene (raw score 0.3719; normalized score 0.039) because it maps to Spinocerebellar ataxia 42 in OMIM (OMIM 616795) (S1 Fig.). The nonsynonymous variant identified in *CACNA1G* (c.1367G>A (NM_018896.4), p.(Arg456Gln)) has a phred-scaled CADD score of 16.13 and is predicted to be deleterious or damaging by several *In Silico* tools (POLYPHEN2, Mutation Taster, FATHMM, Provean, MetaSVM and Meta LR). The top three predicted genes are *PPP2R5B* (intronic variant; normalized score 0.4464), *CASP6* (synonymous variant; normalized score 0.021) and *ADAM11* (synonymous variant; normalized score 0.016).

#### CACNA1G

We evaluated all candidate genes prioritized by phenolyzer in a previously published WES dataset of ET families [15]. We identified one family (S2 Fig.) with a nonsynonymous variant in *CACNA1G* (c.3635G>A (NM_018896.4), p.(Arg1212Gln)), rs150972562) that is highly conserved evolutionarily and is predicted to be deleterious or damaging by several *in silico* tools (provean (score: -3.62), SIFT (score: 0.002) and Mutation Taster (disease causing)) that co-segregated with ET. This *CACNA1G* variant was apparent retrospectively but was not identified in the prior analysis using the bioinformatics pipeline or analysis methods applied in the WES study. The allele frequency of rs150972562 in the Exome Aggregation Consortium (ExAC) database is 0.001596 (192/120264+1 homozygote), which is below the estimates of the disease prevalence of ET at 2-4%.

## DISCUSSION

In this study, we applied the MM-KBAC test [19] to analyze rare variants in the WGS data generated from eight early-onset ET families enrolled in the family study of Essential Tremor (FASET). While numerous methods have been described for rare variant analysis in case-control studies, relatively few methods exist for family-based studies. The advantages of family-based studies are their robustness to population stratification [42], and the use of information about transmission of genetic factors within families, which is more powerful than population-based case control studies [43]. Genes identified by MM-KBAC analysis in ET families were prioritized using phenolyzer. Phenolyzer prioritizes candidate genes based on disease or phenotype information. Phenolyzer includes multiple components, including a tool to map the user-supplied phenotype to related disease, a resource that integrates existing knowledge on disease genes, an algorithm to predict previously unknown disease genes, a machine learning model that integrates multiple features to score and prioritize candidate genes and a network visualization tool to examine gene-gene and gene-disease relationships [39]. Previously, we performed WES [15] on a subset of the families (Families A, B, E and F) included in the current WGS analysis. For these families, WES did not identify the candidate genes identified in the current WGS study. There are several reasons why variants and candidate genes could have been missed in the prior WES analysis. Recently published studies [18, 44] suggest that WGS is more powerful than WES for detecting potential disease-causing mutations within WES regions, particularly those due to SNVs. WGS which forgoes capturing is less sensitive to GC content and is more likely than WES to provide complete coverage of the entire coding region. Other factors that can affect variant and candidate gene identification include the bioinformatics pipeline (GATK version and implementation options) used and statistical analysis methods (WES study pVAAST [15, 45] versus WGS in the current study: MM-KBAC [45]).

In the current study, within each ET family, we generated a prioritized candidate gene list that can be considered for functional studies. In family H, *CACNA1G* is predicted to be the most disease relevant seed gene because it maps to Spinocerebellar ataxia 42 (SCA42) in OMIM (OMIM 616795). *CACNA1G* is also a genetic modifier of epilepsy [46, 47]. The identification of a second family, with a deleterious/damaging *CACNA1G* variant, from a previously published WES dataset strongly suggests that *CACNA1G* may be a susceptibility gene for ET. SCA42 is an autosomal dominant neurologic channelopathy disorder characterized predominantly by gait instability, tremor (i.e. intention, postural, head, and resting) and additional cerebellar signs (i.e. dysarthria, nystagmus and saccadic pursuits), and is caused by a heterozygous mutation in *CACNA1G.* There is variable age at onset (range 9- >78 years) and slow progression of the disease. We reviewed the clinical data in the *CACNA1G* families for the characteristic signs of SCA42 including ataxia, gait instability and ocular signs [48-50]. None of the individuals with ET in these families exhibited these problems, suggesting that these families do not have SCA. On the other hand, neuropathologic studies available for an 83 year old affected individual with SCA42, who also had dementia, showed cerebellar atrophy with Purkinje cell loss and loss of neurons in the inferior olive [48], which in terms of the Purkinje cell loss, is consistent with neuropathological findings of some ET patients [51].

The *CACNA1G* gene encodes the pore forming subunit of T-type Ca(2+) channels, Ca_V_3.1, and is expressed in various motor pathways and may serve different functions [52]. The T-type calcium channel, Cav3.1, has been previously implicated in neuronal autorhythmicity [53, 54] and is thought to underlie tremors seen in Parkinson’s disease[55], enhanced physiological tremor, and in ET [56] and T-type calcium channel antagonists have been shown to reduce tremor in mouse models of ET [54, 57, 58].The identification of *CACNA1G* in two ET families in the current study is consistent with recent reports of mutations in other ion channel genes in other ET families and the concept that the ETs are channelopathies [14, 15]. We previously reported the identification of a mutation in *Kv9.2* (*KCNS2*), that encodes an electrically silent voltage-gated K^+^ channel α subunit, in a family with pure ET [15]. Kv9.2 is highly and selectively expressed in the brain and modulates the activity of Kv2.1 and Kv2.2 channels, which play a major role in membrane excitability and synaptic transmission and is critical for motor control and other neuronal network functions [59]. In two families with atypical ET, mutations were also identified in genes encoding voltage-gated sodium channel alpha subunits. In a family with epilepsy and ET, a disease-segregating mutation p.(Gly1537Ser) in the *SCN4A* gene was identified and functional analyses demonstrated more rapid channel kinetics and altered ion selectivity, which may contribute to the phenotype of tremor and epilepsy in this family [14]. In a four generation Chinese family, with early onset familial episodic pain and ET, a gain-of-function missense mutation p.(Arg225Cys) in *SCN11A* was identified [60]. Collectively, identification of mutations in a T type Ca(2+) channel (*CACNA1G;* two families, this study), a voltage-gated K^+^ channel α subunit (*Kv9.2; KCNS2*, 1 family), and voltage-gated sodium channel alpha subunits (*SCN4A* and *SCN11A*) in ET families (five total to date) is emerging evidence that problems in regulation of membrane excitability and synaptic transmission, which are important more broadly for motor control and other neuronal network functions, could play a role in the pathophysiology of ET. The genetic basis of ET has so far remained elusive. Given the clinical and genetic heterogeneity observed in ET [11-16], evaluation of ion channel genes as candidate genes for ET is warranted.

In family D, *SLIT3* is predicted to be the most disease relevant gene. A disease association with *SLIT3* in OMIM has not been described. The non-synonymous variant identified in *SLIT3* (c.3505G>C, p.(Val1169Leu); rs144799628) is highly conserved evolutionarily, is predicted to be deleterious or damaging by several *in silico* tools and has an allele frequency in the ExAC database of 0.0006407 (72/112370+2 homozygotes), which is below the estimates of the disease prevalence of ET at 2-4%. A disease association of SNPs in the *SLIT3* gene and genetic risk (models: susceptibility, survival and age-at-onset) for Parkinson disease was previously identified in two independent GWAS datasets [61]. Axon guidance pathway molecules are involved in defining precise neuronal network formation during development and in the adult central nervous system play a role in the maintenance and plasticity of neural circuits. The Slit axon guidance molecules and their receptors, known as Robo (Roundabout) serve as a repellent to allow precise axon pathfinding and neuronal migration during development. Three Slit ligands have been identified in vertebrates with spatio-temporal expression patterns in the nervous system as well as in the peripheral tissue and other organs during development. Slit or Robo null gene animal models (*Drosophila* or mouse) show that Slit-Robo interactions act as a repulsive signal to regulate actin dynamics for axon guidance at the midline for commissural, retinal, olfactory, cortical and precerebellar axons [62]. The mechanism by which *SLIT3* contributes to ET may involve early degenerative changes in the years preceding diagnosis and possibly even during brain development (the miswiring hypothesis). In one published study, the candidate gene, *TENM4*, which is a regulator of axon guidance and central myelination, was identified in three ET families [12]. This finding together with the identification of *SLIT3* as a candidate gene in an ET family in the current study suggests that in some instances ET may be a disorder of axon guidance.

In three families, phenolyzer prioritized genes that are associated with hereditary neuropathies (family A, *KARS*, Charcot-Marie-Tooth disease B (OMIM 613641); family B, *KIF5A*, spastic paraplegia 10 with or without peripheral neuropathy (OMIM 604187); and family F, *NTRK1*, hereditary sensory and autonomic neuropathy IV (OMIM 256800). Among the clinical features of CMTRIB with peripheral neuropathy, electrophysiologic studies show motor nerve conduction velocities of 39.5 and 30.6 m/s in the median and ulnar nerves, respectively consistent with an intermediate phenotype between that of demyelinating and axonal CMT [63]. Heterozygous pathogenic mutations in *KIF5A* are also known to cause an axonal CMT subtype [40]. Interestingly, tremor is known to occur in patients with neuropathies although its reported prevalence varies widely [64]. In a case control study that assessed the presence and severity of tremor using the Fahn-Tolosa-Marin Scale, Archimedes spirals and Bain and Findley spiral score, in 43 consecutively recruited patients with inflammatory neuropathies, twenty seven (63%) patients had tremor (posture or action) with a mean age at tremor onset of 57.6 (11.6) years (widely) [64].

In summary, WGS analysis identified candidate genes for ET in 5/8 (62.5%) of the families analyzed. WES analysis of these families in our previously published study failed to identify candidate genes. One drawback to our study is that structural variants (SVs) and copy number variants (CNVs) were not analyzed. However, recent studies suggest that short read Illumina technology for WGS is unable to accurately identify SVs and CNVs and that long read sequencing (PacBio) or other technologies based on nanochannel arrays, such as the Bionano genomics IRYS next generation platform, are needed for accurate detection [65]

The genes and pathways that we have identified can now be prioritized for functional studies to further our understanding of the pathophysiology of ET using cellular and animal models.

## ACKNOWLEDGEMENTS

We thank the patients and families for participating in the study. We also thank and acknowledge the New York Genome Center for performing and generating the WGS data.

## SUPPLEMENTARY INFORMATION

**S1 Fig. Phenolyzer network analysis of WGS gene findings, disease terms and disease types**.

**S2 Fig. Pedigree for family with *CACNA1G* variant (c.3635G>A (NM_018896.4), p.(Arg1212Gln)), identified from a WES dataset**

Supplementary Information accompanies the paper on the PLOS one website.

